# Survey of Public Assay Data: Opportunities and Challenges to Understanding Antimicrobial Resistance

**DOI:** 10.1101/2019.12.13.874909

**Authors:** Akshay Agarwal, Gowri Nayar, James Kaufman

## Abstract

**ABSTRACT:** Computational learning methods allow researchers to make predictions, draw inferences, and automate generation of mathematical models. These models are crucial to solving real world problems, such as antimicrobial resistance, pathogen detection, and protein evolution. Machine learning methods depend upon ground truth data to achieve specificity and sensitivity. Since the data is limited in this case, as we will show during the course of this paper, and as the size of available data increases super-linearly, it is of paramount importance to understand the distribution of ground truth data and the analyses it is suited and where it may have limitations that bias downstream learning methods. In this paper, we focus on training data required to model antimicrobial resistance (AR). We report an analysis of bacterial biochemical assay data associated with whole genome sequencing (WGS) from the National Center for Biotechnology Information (NCBI), and discuss important implications when making use of assay data, utilizing genetic features as training data for machine learning models. Complete discussion of machine learning model implementation is outside the scope of this paper and the subject to a later publication.

The antimicrobial assay data was obtained from NCBI BioSample, which contains descriptive information about the physical biological specimen from which experimental data is obtained and the results of those experiments themselves.[1] Assay data includes minimum inhibitory concentrations (MIC) of antibiotics, links to associated microbial WGS data, and treatment of a particular microorganism with antibiotics.

We observe that there is minimal microbial data available for many antibiotics and for targeted taxonomic groups. The antibiotics with the highest number of assays have less than 1500 measurements each. Corresponding bias in available assays makes machine learning problematic for some important microbes and for building more advanced models that can work across microbial genera. In this study we focus, therefore, on the antibiotic with most assay data (tetracycline) and the corresponding genus with the most available sequence (*Acinetobacter* with 14000 measurements across 49 antibiotic compounds). Using this data for training and testing, we observed contradictions in the distribution of assay outcomes and report methods to identify and resolve such conflicts. Per antibiotic, we find that there can be up to 30% of (resolvable) conflicting measurements. As more data becomes available, automated training data curation will be an important part of creating useful machine learning models to predict antibiotic resistance.

**CCS CONCEPTS:** • Applied computing → Computational biology; Computational genomics; Bioinformatics;

## 1 INTRODUCTION

Introduction of machine learning and other computational techniques to the field of biology and medicine have revolutionized the way research can be conducted in these disciplines[2]. Due to machine learning, researchers are now able to leverage the power of data in order to identify patterns that can potentially help solve important problems, such as antimicrobial resistance (AR) [3–9]and detecting food hazards [10–14]. Computational advances in these areas has also led to the emergence of consumer-centric industries. Companies like uBiomeTM provide information based on nucleotide sequence data and computational biology, about potential health risk, physical characteristics, and ancestry[15]. Since this analysis is used for real-world health decisions, we must ensure the predictions are meaningful and accurate.

However, achieving reliable predictions require a considerable amount of curated ground truth data [16, 17]. There are several projects to categorize reference lists of genes associated with AR, each with different curation criteria[18–20]. Collectively, AR genes are often described as part of a “resistome” [21]. A number of issues confound the ability to collect reliable training data for AR prediction. There are multiple mechanisms of resistance to antimicrobial compounds (drugs). Resistance mechanisms vary by compound. Observation of an individual resistome, while correlated with resistance to a compound, is not always a predictor of resistance for that compound[18, 19]. Furthermore the phenotypes, resistant or susceptible, are not boolean properties. Instead, AR should be understood as a stress response that results in non-discrete phenotypes. Antimicrobial compounds subject microorganisms to a stress, the nature and magnitude depending on the dosage and type of compound. Microorganisms respond to this stress by a number of mechanisms which include, but are not limited to:

- enzymatic deactivation of the drug
- alteration of the drug target- or protein binding site
- changes to metabolic pathways
- reduced cell wall permeability to reduce absorption of the compound
- or activation of efflux to pump the drug out of the cell[5, 18].

Many current approaches for resistance prediction perform phenotype classification i.e. for a given antibiotic-accession pair, determine whether the bacteria will be resistant, susceptible, or intermediate to the antibiotic. To achieve this, they rely upon methods akin to gene counting. In gene counting, a set of genes believed to cause resistance to an antibiotic are counted, after annotating the genome. A heuristic based on presence of these genes is used to predict the phenotype.

For the above approach to be successful, the list of known resistant genes must be continuously updated and the requirement of resistant genes co-occurring may get discounted. Also, single nucleotide polymorphism (SNP) changes can become difficult to predict. As new resistant genes are discovered, the above approach may fail or provide inadequate results.

AR response varies by organism and is based on an organism’s particular resistome. However, as a part of the stress response, microbes can acquire new genes through transfer of plasmids, integrative conjugative elements, or other horizontal gene transfer (HGT) mechanisms [22, 23]. As additional stress response pathways are activated, the organism can survive an increased concentration of the antimicrobial compound. So the organisms ability to ‘resist’ a treatment is a function of dose, or concentration. A clinician prescribes antimicrobial medication based on an optimal or maximal safe dosage, but that dosage may be different if the patient is a human or a large livestock animal. Clinical laboratories use one of several tests to measure the ability of an organism to grow (or to survive) as a function of dosage. One widely used test provides a laboratory estimate of the *minimum inhibitory concentration* or MIC[24]. Data from tests like MIC are used to select secondary treatments for problematic infections. Any machine learning or AI approach to predicting resistance should, therefore, focus on predicting MIC. The phenotype, or prediction, of an organism’s response to antimicrobial treatment can’t be decoupled from the prediction of MIC, since the predicted phenotype depends on the compound, the infectious organism genome, and the patient governing the maximal safe concentration for the therapy.

The CDC terms AR as a One Health problem [25] i.e. the health of people is related to the animals and environment, since 6 out of 10 infectious diseases spread to humans from animals. MIC prediction helps tackle the problem at multiple levels, such as food production, food safety, and animal health.

In this paper, we describe both the challenges and opportunities of using biological assay data available through public repositories, namely NCBI’s BioSample [26]. We begin by introducing the important terms, tools and technologies that we used/using during the course of this paper.

1. NCBI - National Center for Biotechnological Information, acts a central repository for reporting results, data, research and tools.[20]
2. BioSample - Database containing biological aspects of sequencing experiments./citebarrett2011bioproject
3. Antibiogram - Data about antimicrobial susceptibility and resistance derived from drug resistant pathogens submitted to BioSample.[27]
4. Assay[biochemical] - A biochemical assay is an analytical in-vitro procedure used to detect, quantify and/or study the binding or activity of a biological molecule, such as an enzyme. [28]
5. MIC - minimum inhibitory concentration - lowest concentration of a chemical that inhibits visible bacterial growth.[28]

We will then discuss the distribution of the data to highlight the analysis the data is suited for, and the challenges in using this data as ground truth for machine learning tasks. Furthermore, we discuss an approach to clean this data, identify, and handle conflicts for identifying ground truth MIC.

## 2 DATA

### 2.1 Raw Data and Transformations

From NCBI BioSample, we gathered the bacterial assay data, resulting in 78000 assays. The headers mined from the XML file are:

- COMPOUND
- SRA_ID
- BIOSAMPLE_ACCESSION
- PHENOTYPE_DESCRIPTION
- MEASUREMENT
- MEASUREMENT_SIGN
- MEASUREMENT_UNIT
- TYPING_METHOD
- TYPING_PLATFORM
- VENDOR
- LAB_TYPING_METHOD_VERSION
- TESTING_STANDARD

This raw data, hereto referred to as Data Stage 1, included 99 different antibiotic compounds associated with 4962 SRA_IDs (each SRA_ID indicates a bacterial whole genome isolate sequence data set) and 5173 BioSample accessions (Table 1). “Phenotype description” relays the observable physical trait resulting from the antibiotic assay which is comprised of 6 potential values: resistant, susceptible, intermediate, not defined, susceptible-dose dependent, and non-susceptible. There are 5 assay typing methods, 15 typing platforms, 12 vendors, 16 lab typing method versions and 6 testing standards such as MIC, CLSI, and agar dilution. Thus, the amount of variations for each assay greatly reduces the number of reference data points per class. Table 2 lists, in particular, the count of *MIC Assays* per compound.

**Table 1:** Assay Distribution - Data Processing.

| Data Stage | #Antibiotics | #SRA_ID | #Genomes | #assays |
| --- | --- | --- | --- | --- |
| Data Stage 1 | 99 | 4963 | 5173 | 77424 |
| Data Stage 2 | 74 | 4962 | 4962 | 73600 |
| Data Stage 3 | 50 | 1399 | 1399 | 30076 |

Note: the compounds associated with only a single assay were removed as there was insufficient data. The final distribution of ground truth data after conflict resolution is listed in Table 3 and Table 4.

**Table 2:** Assay Distribution by Antibiotic.

| Antibiotic | # MIC Assays |
| --- | --- |
| ciprofloxacin | 3542 |
| gentamicin | 3355 |
| trimethoprim-sulfamethoxazole | 3342 |
| tetracycline | 3224 |
| ceftriaxone | 3122 |
| ampicillin | 2873 |
| cefoxitin | 2638 |
| amikacin | 2243 |
| amoxicillin-clavulanic acid | 2138 |
| levofloxacin | 1826 |
| ceftazidime | 1795 |
| meropenem | 1710 |
| imipenem | 1663 |
| chloramphenicol | 1603 |
| tobramycin | 1581 |
| kanamycin | 1466 |
| cefotaxime | 1433 |
| azithromycin | 1425 |
| nalidixic acid | 1422 |
| sulfisoxazole | 1420 |
| ceftiofur | 1420 |
| streptomycin | 1420 |
| cefazolin | 1352 |
| aztreonam | 1320 |
| ampicillin-sulbactam | 1185 |
| cefepime | 1119 |
| piperacillin-tazobactam | 1046 |

In order to predict MIC values for a given antibiotic, we process and label data according to the following defined stages:

1. raw data (Data Stage 1).
2. removal of all rows missing values for the 5 headers shown in italics above (Data Stage 2).
3. To ensure self consistency, we use data only for high quality, complete genomes. For these data sets, we downloaded the sequence data, assembled it into near-complete genomes, and annotated them in order to identify all relevant nucleotide and amino acid sequences (Data Stage 3).

Details about the distribution of data at each stage is shown in Table 1.

We were left with 30076 assays, with 50 antibiotics and 1399 accessions across 19 genera.

Note: the compounds associated with only a single assay were removed as there was insufficient data. The final distribution of ground truth data after conflict resolution is listed in Table 3 and Table 4.

**Table 3:** Data Point Distribution by Antibiotic - Data Stage 3.

| Antibiotic | # of MIC Data Points | Antibiotic | # of MIC Data Points |
| --- | --- | --- | --- |
| trimethoprim-sulfamethoxazole | 1336 | penicillin | 146 |
| ciprofloxacin | 1314 | clindamycin | 145 |
| tetracycline | 1296 | chloramphenicol | 142 |
| ceftriaxone | 1240 | cefotetan | 118 |
| gentamicin | 1234 | moxifloxacin | 97 |
| levofloxacin | 1204 | colistin | 97 |
| amikacin | 1147 | polymyxin B | 91 |
| imipenem | 1125 | cefuroxime | 81 |
| tobramycin | 1111 | vancomycin | 74 |
| ceftazidime | 1107 | rifampin | 73 |
| ampicillin | 1105 | linezolid | 73 |
| cefotaxime | 1067 | nalidixic acid | 72 |
| aztreonam | 1026 | ceftiofur | 71 |
| cefazolin | 1011 | streptomycin | 71 |
| cefoxitin | 881 | synercid | 71 |
| ampicillin-sulbactam | 871 | sulfisoxazole | 71 |
| meropenem | 844 | amoxicillin | 71 |
| nitrofurantoin | 725 | azithromycin | 71 |
| doripenem | 506 | kanamycin | 70 |
| tigecycline | 498 | minocycline | 56 |
| cefepime | 418 | ticarcillin-clavulanic acid | 16 |
| piperacillin-tazobactam | 401 | doxycycline | 13 |
| amoxicillin-clavulanic acid | 390 | cefotaxime-clavulanic acid | 9 |
| ertapenem | 332 | fosfomycin | 2 |
| erythromycin | 146 | daptomycin | 2 |

**Table 4:** MIC Data Point Dist by Genus - Data Stage 3.

| Genus | # |
| --- | --- |
| *Acinetobacter* | 13992 |
| *Klebsiella* | 4423 |
| *Streptococcus* | 1567 |
| *Escherichia* | 1456 |
| *Salmonella* | 1109 |
| *Enterobacter* | 839 |
| *Pseudomonas* | 322 |
| *Serratia* | 178 |
| *Citrobacter* | 69 |
| *Proteus* | 46 |
| *Staphylococcus* | 29 |
| *Kluyvera* | 23 |
| *Morganella* | 23 |
| *Providencia* | 23 |
| *Shigella* | 21 |
| *Achromobacter* | 12 |
| *Corynebacterium* | 4 |
| *Listeria* | 2 |

Table 3 refers to all genome accessions independent of microbial genus. Table 4 shows how the curated measurements are distributed across genera.

### 2.2 NCBI Sample Bias

From Table 4 significant bias is evident in the NCBI Sequence data where sequence and assay coverage is more complete for some genera than others. This in part reflects the current focus of public funding and pathogen occurrence, as well as emerging resistance in important clinical organisms. Since the set of resistance mechanisms can vary by organism, this bias impedes balanced training data across mechanisms and genera.

## 3 METHODS

### 3.1 Data Download and Extraction

The complete NCBI BioSample data, biosample_set.xml.gz, was downloaded from the NCBI ftp server. SRA_Run_Members.tab and SRA_Accessions.tab files were downloaded from metadata reports from the NCBI ftp server. BioSample metadata is an xml file that contains multiple fields relating the group/organization that conducted the experiment, GIS data, source of the isolate, as well as the Antibiogram data. The SRA_Run_Members.tab file contains mappings between BioSample accessions, run accessions, experiment IDs, and sequencing run IDs.

BioSample data was parsed using an XML SAX parser, built in Java, and the bacterial Antibiogram data was extracted. We parsed the SRA_Run_Members.tab and extracted the SRA_ID and BioSample_Accession from the same source. The data was further analyzed using Python scripts.

#### Genome Assembly and Annotation

All bioinformatics tools subsequently described are open source. To assemble bacterial sequence isolates retrieved from the NCBI SRA, Trimmomatic 0.36 [29] was used to remove poor-quality base calls, poor-quality reads and adapters from the sequence files. For removal of PhiX control reads, Bowtie 2 2.3.4.2 [30] was used to align the sequences to references derived from PhiX174 (Enterobacteria phage phiX174 sensu lato complete genome). FLASh 1.2.11 [31] was used to merge paired-end reads from the resulting sequences to improve quality of the assembly. Once these quality control steps were completed, the merged reads were assembled using SPAdes 3.12.0 [32] and QUAST 5.0.0 [33] in an iterative assembly/quality evaluation process. After genome assembly, genes and proteins were annotated using Prokka 1.12. [34] For 1399 high quality complete genomes with antibiogram data, this annotation process identified all unique gene and protein sequences.

#### Genome Curation and Selection

The accuracy of metadata and quality of WGS (wholge genome sequence) files maintained by NCBI varies dramatically. Some files, for example, may have been derived from wet lab contaminated bacterial isolates (non-pure culture containing more than one microbe) and are therefore not representative of the single isolate labeled. Others may be labeled incorrectly with a mismatched microbial genus ID, likely a result of human clerical error. To optimize for sufficient sequencing depth, only bacterial SRA datasets at least 100 MB in size were downloaded and converted to FASTQ format using SRA-toolkit [35]. To maintain a high-quality data set of more complete assemblies, genome assemblies containing greater than 150 contigs of size > 500 bp (base pairs) and an N50 of less than 100,000 bp were discarded, with the exception of genomes from the genus *Shigella*. For *Shigella*, assemblies containing greater than 500 contigs (of size > 500 bp) and an N50 of less than 15,000 bp were discarded.

### 3.2 Extracting Features from Assays

Biochemical assays include measurements for a particular compound (e.g., antibiotic) at various concentrations tested with a specific cultured organism. For studies of AR, as stated each data point in an assay will have an outcome associated with a particular concentration reflected in "Phenotype description". Resistance at a particular concentration does not necessary imply the organism is resistant to the antibiotic at all concentrations. Here, susceptible implies the organism does not grow (or, for some protocols, that it dies) at a particular concentration. At very low concentrations, resistance is the expected outcome for any compound. To classify the organism’s susceptibility one needs to know the minimum inhibitory concentration (MIC), defined by the maximum concentration at which the organism is resistant and/or the minimum concentration at which it is susceptible. It is then necessary to compare the critical MIC obtained from the assay to the maximum safe does for the patient, which varies between humans and livestock. Different laboratories use different protocols in conducting assays. Some labs may stop measurements if, for example, the organisms is found to be susceptible at low concentration or resistant above a defined concentration.

In this study, we first categorized all assays by specific antibiotic and then by genome accession. For each antibiotic-accession pair we then created a resistant values list, a susceptible values list and an intermediate values list. Each list contained only two entries, min concentration and max concentration for each category. Thus, for each antibiotic-accession pair, we extracted from the assay unique the resistant_min, resistant_max, susceptible_min, susceptible_max, intermediate_min, and intermediate_max.

In order to do the above, we parse each assay and identify the antibiotic-accession pair. Then we look at the phenotype description - resistant, susceptible or intermediate. This dictates which list, resistant, intermediate or susceptible, to update. We then look at the measurement and the measurement sign and accordingly update the min or max values of the relevant list.

The data from some assays may contain only a range of resistant or a range of susceptible concentrations, thus a switch from resistant to susceptible is not observed. Therefore, for each assay we extract the two important values, i.e. min susceptible and/or max resistant concentration, either of which can define the MIC value described above. However, it is still necessary to resolve experimental error, noise, and associated conflict within this data.

In order to further clean the data we have obtained here, we define and describe the steps in the next two sections, Conflict Identification and Conflict Resolution.

### 3.3 Conflict Identification

One of the most important factors to consider while selecting input data for a machine learning task is data cleanliness. Noisy data can often produce spurious results and degrade performance, even if the evaluation scores are high.

In assay data, conflicts are a significant problem due to the scarcity of the data. Consider a subset of fields from Antibiogram, namely, SRA_ID (genome accession), phenotype_description, measurement_sign, measurement and compound (antibiotic). From this data we identify two types of conflicts:

- Direct Conflict: where all the fields except phenotype_description are identical
- Range Conflict: where resistant, intermediate and susceptible ranges overlap with one another as shown in Figure 3.

Up to 10% of measurements for each genus were identified as conflicts, with the largest number of conflicts, 1441 values, within *Acinetobacter*. Up to 35% of measurements for some antibiotics were conflicting, with ceftriaxone contaning the most number of conflicts, 434 values. [Note: For sake of brevity, we haven’t included full table on number of conflicting entries.]

There can be multiple reasons for conflicts in phenotypes, including differences in testing standards or testing equipment. However, irrespective of these differences, it is necessary to clean the data and ensure that confounding conflicts are not passed on to the machine learning model. It is also important to note that we measure conflicts for a particular antibiotic-accession pair, thus evolution of genomes need not be considered here as they will be captured by different accessions.

We have thus come up with a novel method to transform the data such that these conflicts are resolved and do not impede model’s learning rate. Note: Resolved means extraction of probable MIC value.

### 3.4 Conflict Resolution

To curate the assay data we developed approaches to resolve both the range conflicts and direct conflicts discussed above. These methods can potentially be improved through future work by looking at additional fields from Antibiogram including typing_method, typing_version, vendor, testing_standard etc. One may further look at metadata from Biosample in order to strengthen curation and isolate noise.

#### 3.4.1 Range Conflicts

Figure 3 provides a visual example of a range conflict. Over a range of concentration, multiple data points are associated with conflicting outcomes. In this example, some concentrations are labeled both resistant and intermediate or both susceptible and intermediate. Here, the conflict can be conservatively resolved by adopting the higher susceptible concentration and lower resistance concentration, to avoid erroneously predicting the drug is effective when there is a discrepancy.

Figure 4 describes the approach to range conflict resolution. In order to decide which range to resize, we use the confidence value. The underlying aim is to ensure that the MIC value is correctly identified. To define MIC value, we use the higher value between the maximum resistant value and the minimum susceptible value. Thus, we need to resolve each range and rely on the following: *MIC = max {minimum_resistant,maximum_resistant, minimum_intermediate,maximum_intermediate, minimum_susceptible}*

We include both minimum_resistant and maximum_resistant as well as both minimum_intermediate and maximum_intermediate because in most cases only one the above values is obtained from the assays.

#### 3.4.2 Direct Conflicts

To resolve direct conflicts, we check if the measurement value and phenotype agree with the new ranges. If there is agreement, we keep the assay and increase the confidence value, else we discard it.

In the Figure 1 and 2 we look at the list of antibiotic-accession pairs and the associated min/max values in the resistant,susceptible and intermediate categories. For each data point we plotted the concentration, min and max, and colored them based on phenotype. However, both the min and max value existed in each phenotype category for only a few antibiotic-accession pairs.

**Figure 1:**
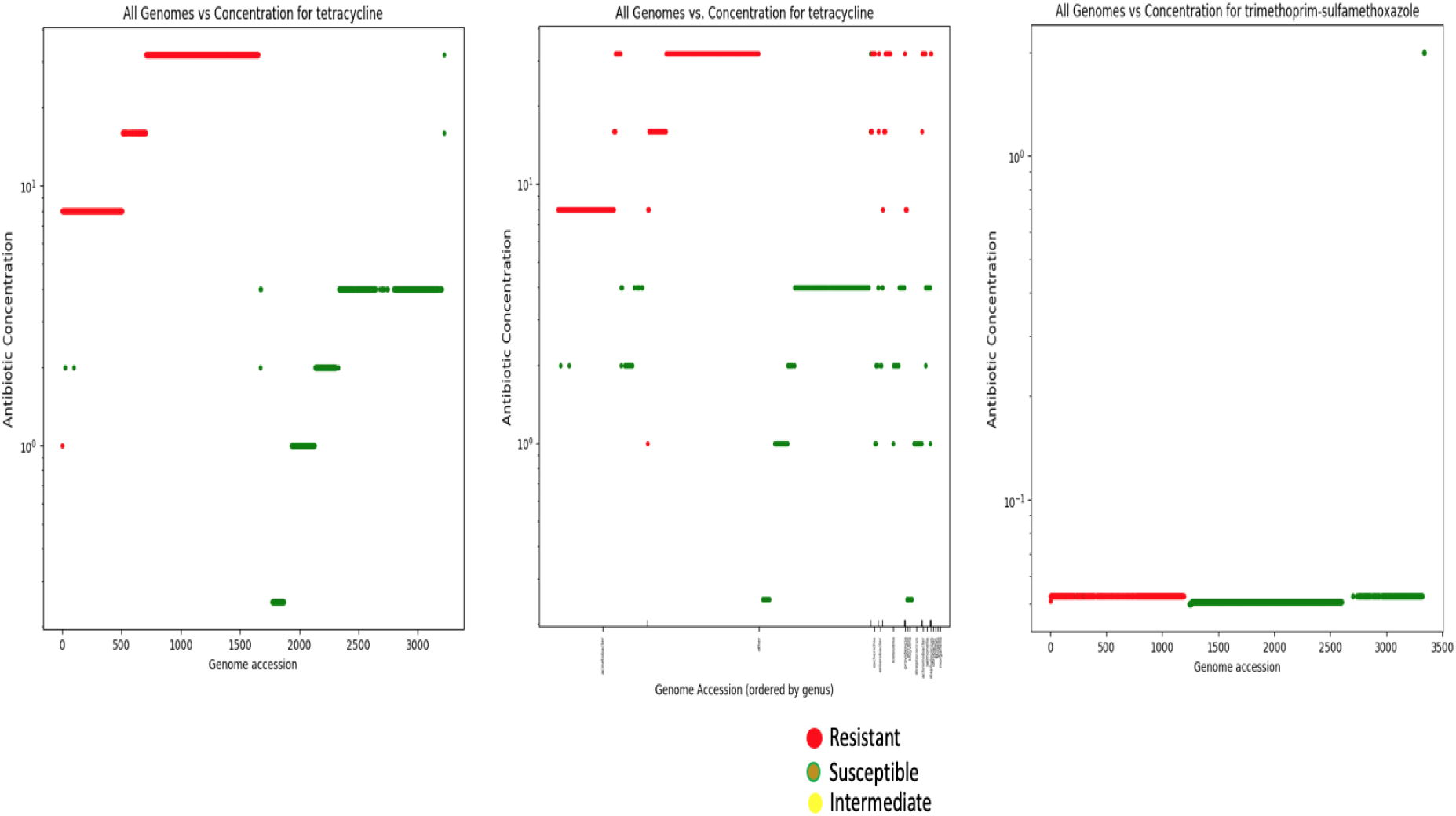
MIC Assay Distribution - Tetracycline,Trimethoprim-Sulfamethoxazole.

**Figure 2:**
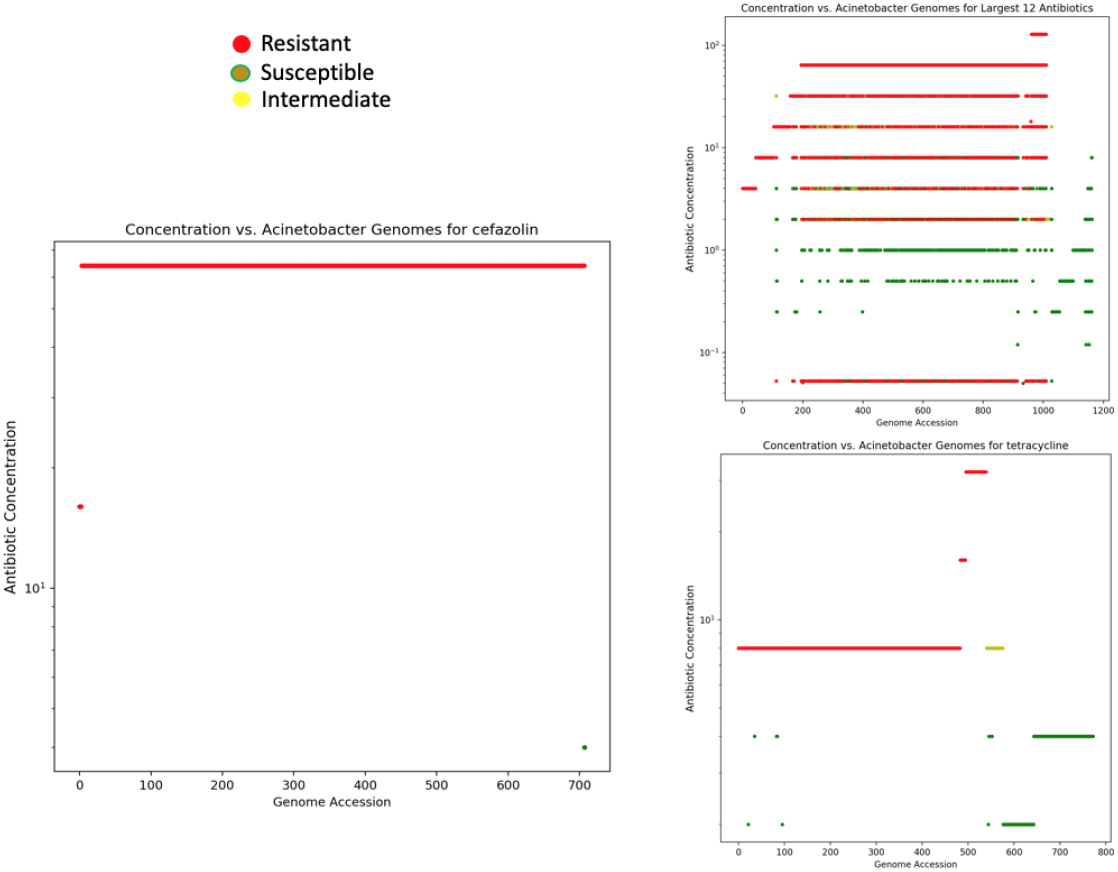
MIC Assay Distribution - *Acinetobacter* with Cefazolin and Tetracycline.

**Figure 3:**
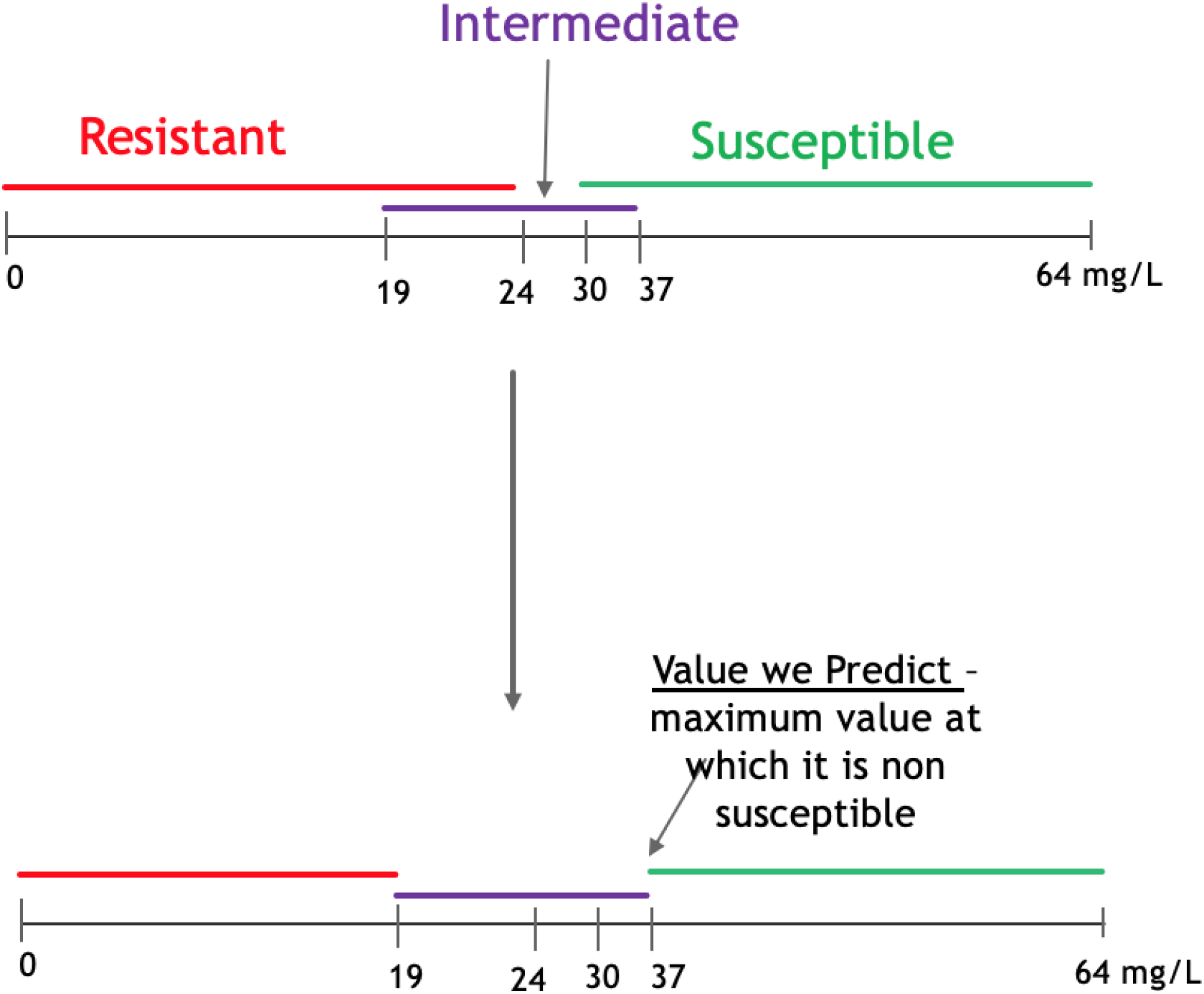
A visual example of a range conflict. Multiple data points are associated with conflicting outcomes. In the example, some concentrations are labeled both resistant and intermediate, or susceptible and intermediate.

**Figure 4:**
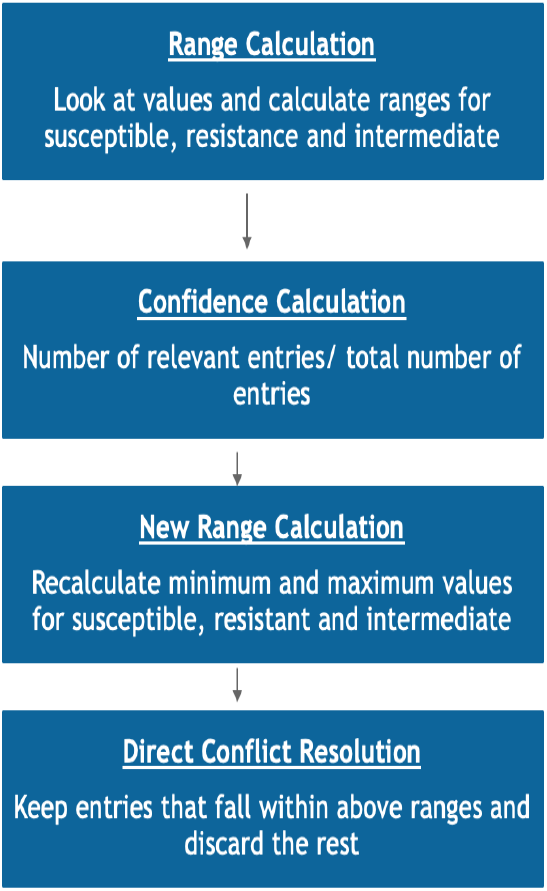
Conflict Identification and Resolution.

## 4 DISCUSSION

### 4.1 Evaluation of Ground Truth Data

#### Availability and Biases

We now look at the distribution of Data Stage 2 which is a result of the transformation where we removed all assays with null SRA_ID and testing standard other than MIC (see Section 2 Data).

To analyze Data Stage 2, we plotted the distribution of min susceptible and max resistant MIC values for each antibiotic after sorting them by phenotype description. We placed the assays with resistant phenotype first, followed by intermediate phenotype, and then susceptible. We did not consider values where the MIC was listed as ‘not defined‘, ‘susceptible-dose dependent’ or ‘non-susceptible‘. The number of assays with these ambiguous phenotype descriptions was minimal from the complete set of 73600 measurements. The MIC distribution for Tetracycline has been shown in Figure 1.

In Figure 1,in the left most visualization, the x-axis is the genome accessions and y-axis is the concentration value. We see that resistant values are at higher concentrations and susceptible values are found to be at lower concentrations, which may seem contrary to expected observations. This could reflect underlying lab procedures. If an assay reveals resistance at concentrations higher than approved maximal dose, the lab does not continue to test for susceptibility at even higher concentrations since the drug can not be prescribed at that dose. Conversely, isolates found to be susceptible at concentrations at or below approved dosages need not be tested for resistance at much lower dosages. The divide between red and green reflects a concentration range determined by approved clinical practice. From a machine learning perspective, depending on the model and use case, it can be important to pass ranges instead of singular values.

We can use such data for a phenotype classifier, with input features as representation of the genomes, such as component genes, along the x-axis. From the distribution of the cleaned training data, we can expect the classifier to perform reasonable well. However, using the same data to perform a regression task poses major problems, since it is difficult to fit a regression curve to the distribution. Thus, such analysis highlights both the type of model it is suited for, as well as they type of prediction possible, phenotype rather than MIC value.

In Figure 1, the graph on the right shows the distribution for Trimethoprim-Sulfamethoxazole. We observe a significant number of data points at similar values.Since most assays are of *Acinetobacter* genomes, the high level of homogenity between the features (gene sequences) and lack of data makes construction of a sensitive model, for both phenotype classification and MIC prediction, difficult.

Figure 2 shows the distribution of acinetobacter assays for tetracycline and cefazolin. An interesting point to note here is that, for cefazolin, we have only one assay for susceptible phenotype. This indicates that it will be difficult to train a model for cefazolin with acinetobacter.

### 4.2 Data Stage 3

For Data Stage 3, we created a list of high quality genomes from the subset of SRA ids with assay data. We then proceeded to download the raw sequence data from NCBI, assembled it and used Prokka in order to annotate these genomes and get a list of nucleotide and amino acid sequences contained within these genomes.[34]

From Table 3, we see that once we subset the assays to include only those with high quality and complete genomes, the data is considerably reduced per antibiotic.

From Table 4, we see that the largest number of assays are present for the genus *Acinetobacter* and the least is for *Listeria*. This indicates another possible bias in the sequencing data and reflects that *Acinetobacter* poses a more immediate threat in areas like antimicrobial resistance.

### 4.3 AntiMicrobial Resistance Prediction

We have discussed data distribution, data cleaning, identification of conflicts in the assay data and a simple, effective method to handle these conflicts. As we had remarked earlier, this cleaned data is used for machine learning tasks, specifically tasks like prediction of antimicrobial resistance.

In order to perform AR prediction, it is crucial to determine which entity to predict, instead of simply predicting the resistance phenotype, resistant, susceptible or intermediate, for a genome.

As described previously, MIC is the highest concentration at which bacteria is resistant to the antibiotic or the lowest concetration at which it is susceptible.

Furthermore, in order to predict the resistance phenotype, one can simply identify thresholds of safe concentrations of antibiotics for different target users and thus assign a phenotype based on the minimum safe concentration predicted.

Though the discussion of a machine learning model implementation is outside the scope of discussion for this paper, we would like to briefly mention our results from the AR MIC prediction model, as it serves to highlight the idea of how to use this assay data as training input to a model. Using XGBoost we were able to achieve an R squared value of 0.67 and for phenotype classification, we were able to achieve an accuracy of 0.94 using Decision Trees for Tetracycline. The feature vector consisted of component genes sequences of genomes overlapping with known resistant gene sequences found in Megares [18]. We are working on improving our approach, however the high accuracy of phenotype classification derived from MIC prediction is a promising result and more robust than techniques like gene counting.

Many machine learning models are showing significant promise in prediction of both resistant genes[36] and mic[3].

### 4.4 Partitioning the Data

Since different antibiotics target different cellular processes, and are subject to different resistance mechanisms, the gene features relevant to prediction of resistance vary by drug or compound. For this reason, it is necessary to partition the ground truth data by compound. In addition, drugs are often prescribed based on organism name or genus. In some sense, partitioning by genus is a surrogate for our incomplete understanding of the potential set of genes that contribute to resistance. Since the core genomes differs by genus, partitioning training data by genus may reduce conflicts, but reflects an incomplete understanding of the full resistome. E.g. Chimeric genomes are on the border of multiple genera. Resistance genes are often found on plasmids and/or integrative conjugative elements.

### 4.5 Insufficient Data

From the table 3 and 4, we observe that there is a serious lack of data to perform effective MIC prediction or even phenotype classification. The lack of ground truth data points across the entire spectrum of concentration levels is not conducive to a linear regression model. Furthermore a more complete coverage of data would allow for more specificity when identifying the gene features contributing to resistance to a particular antibiotic. Thus, the current data set calls for development of robust mathematical models relying on minimal training data to create accurate predictions for use cases in AR.

## 5 CONCLUSIONS

We have arrived at the following conclusions:

1. The existing data contains conflicts which need to be resolved before the data in passed as input to machine learning models to ensure higher sensitivity and specificity.
2. We outline a method for conflict identification and for extraction of MIC values that can be used to train AR/MIC prediction models.
3. There is a serious lack of data per antibiotic and per genus. For even the largest set for *Acinetobacter* is not enough to train an effective machine learning model.
4. Data distribution makes it harder to train regression models even though classification models may be easier to train.
5. It is important to be cognizant of the biases in data e.g. Figure 2 while developing mathematical/computational models.
6. It is important to focus on developing robust mathematical models that can create accurate predictions even with little data. 5. On the basis of existing data and using tools like BLAST can we approximate more data? this can be appended to our machine learning model results.

## 6 FUTURE WORK

Current analysis can be extended in the future in multiple ways, some of them being:

1. Analysis of metadata such as source, lab typing method etc in order to isolate more concretely noisy or outdated assays.
2. Exploration into ways to extend the data set using artificial techniques and surrogate models.

## ACKNOWLEDGMENTS

The authors would like to thank Dr. Kristen Beck and Dr. Vandana Mukherjee for helpful discussion and improvements to the manuscript.

